# Self-reported health behaviors and longitudinal cognitive performance: Results from the Wisconsin Registry for Alzheimer’s Prevention

**DOI:** 10.1101/742700

**Authors:** Kimberly D. Mueller, Rebecca L. Koscik, Derek Norton, Martha C. Morris, Erin M. Jonaitis, Lindsay R. Clark, Taylor Fields, Samantha Allison, Sara Berman, Sarah Kraning, Megan Zuelsdorff, Ozioma Okonkwo, Nathaniel Chin, Cynthia Carlsson, Barbara B. Bendlin, Bruce Hermann, Sterling C. Johnson

## Abstract

**Background:** Studies have suggested associations between self-reported engagement in health behaviors and reduced risk of cognitive decline. Most studies explore these relationships using one health behavior, often cross-sectionally or with dementia as the outcome. In this study, we explored whether several individual self-reported health behaviors were associated with cognitive decline when considered simultaneously, using data from the Wisconsin Registry for Alzheimer’s Prevention (WRAP), an Alzheimer’s disease risk-enriched cohort who were non-demented and in late midlife at baseline.

**Method:** We analyzed longitudinal cognitive data from 828 participants in WRAP, with a mean age at baseline cognitive assessment of 57 (range = 36-78, sd = 6.8) and an average of 6.3 years (standard deviation = 1.9, range = 2-10) of follow-up. The primary outcome was a multi-domain cognitive composite, and secondary outcomes were immediate/delayed memory and executive function composites. Predictors of interest were self-reported measures of physical activity, cognitive activity, adherence to a Mediterranean-style diet (MIND), and interactions with each other and age. We conducted linear mixed effects analyses within an Information-theoretic (IT) model averaging (MA) approach on a set of models including covariates and combinations of these 2- and 3-way interactions. The IT approach was selected due to the large number of interactions of interest and to avoid pitfalls of traditional model selection approaches.

**Results:** Model-averaged results identified no significant modifiable behavior*age interactions in relationship to the primary composite outcome. In secondary outcomes, higher MIND diet scores associated with slower decline in executive function. Men showed faster decline than women on delayed memory, independent of health behaviors. There were no other significant interactions among any other health behaviors and cognitive trajectories.

**Conclusions:** When multiple covariates and health behaviors were considered simultaneously, there were limited weak associations with cognitive decline in this age range. These results may be explained alone or in combination by three alternative explanations: 1) the range of cognitive decline is in middle age is too small to observe relationships with health behaviors, 2) the putative associations of these health behaviors on cognition may not be robust in this age range, or 3) the self-reported measures of the health behaviors may not be optimal for predicting cognitive decline. More study may be needed that incorporates sensitive measures of health behaviors, AD biomarker profiles, and/or other disease comorbidities.

## Introduction

The estimated prevalence of dementia in America in 2019 was 5.8 million, and it is projected to rise to 13.9 million by 2050 if prevention or cures are not identified. Delaying the onset of dementia by five years would result in a financial savings of up to 40% in 2050^1^, as well as a reduction of emotional burden to families and caregivers. Consequently, there is considerable interest in identifying lifestyle modifications that may prevent or delay cognitive decline in aging adults, particularly those with elevated dementia risk.

Numerous observational studies have shown correlations between lifestyle activities, such as engagement in physical, cognitive, and social activities, and cognitive decline in aging adults^2,3^. Furthermore, in the last decade there have been multiple studies showing that consuming a Mediterranean-style diet, or aspects of this type of eating (e.g., fish), also confer a reduced risk of cognitive decline in aging adults^4–6^. Studies that examine multiple health behaviors and cognitive decline from late middle age – the time at which lifestyle interventions may be most effective – are lacking.

Although analyses using lifestyle modifications as predictors often control for factors that may influence cognition such as age, educational attainment, and intellectual ability, studies often include one main predictor of interest (e.g., cognitive activity), but not other lifestyle activities that may tend to cooccur^7^. Such studies do not explain to what degree one lifestyle behavior contributes to variability in cognition above and beyond other health behaviors.

Moreover, advice that targets single health behaviors may have unintended consequences for modifying other health behaviors that tend to co-occur (e.g., increasing physical activity but engaging in poor diet)^8^. The purpose of the present study was to determine whether self-reported health behaviors—cognitive activity, physical activity and diet—were associated with attenuated cognitive decline when considered simultaneously, using data from the Wisconsin Registry for Alzheimer’s Prevention (WRAP).

## Methods

### Participants

WRAP is a longitudinal observational cohort enriched for a parental history of late-onset AD. Enrollment began in 2001 with follow-up assessments occurring every two years; for additional protocol details, see Johnson et al., 2018.^9^ Due to this targeted enrollment, participants have a higher percentage of the *APOE-ε4* allele than the general population (~41%). At the time of these analyses, 1549 participants were enrolled, with a mean age at baseline of 53.7 (sd=6.6), 72.6% had a parental history of sporadic AD, and all were free of dementia at baseline. To be included in these analyses, participants had to have at least two visits; be free at baseline of clinical Mild Cognitive Impairment (MCI) and histories of stroke, Parkinson’s disease, multiple sclerosis or epilepsy at any visit; and had to have complete data for the predictors (completed questionnaires for diet, physical activity, and cognitive activities (n=828)). This study was conducted in compliance with the ethical principles for human subjects research defined in the Declaration of Helsinki, including approval by the University of Wisconsin – Madison Institutional Review Board.

### Cognitive Outcomes

The full WRAP battery is described in greater detail in Johnson et al. (2018)^9^. For the primary outcome, we calculated a composite analogous to the Preclinical Alzheimer’s Cognitive Composite 4 (PACC4) described by Donohue et al. (2014)^10^ based on our available test measures. Specifically, we computed a PACC4 composite as the average of standardized test scores (z-scores) of total recall learning trials 1-5 from the Rey Auditory Verbal Learning Test (RAVLT)^11^, total scores for the Logical Memory II subtest (i.e., delayed recall of stories A and B) from the Wechsler Memory Scale—Revised (WMS-R)^12^, total scores from the Digit Substitution test of the Wechsler Abbreviated Intelligence Scale-Revised (WAIS-R)^13^, and the total score from the Mini-Mental Status Examination (MMSE)^14^.

#### Secondary outcomes

We also averaged standardized test scores to obtain three domain-specific cognitive composites previously described by our group^15^, including Immediate Recall (contributing tests: RAVLT total from trials 1-5; WMS-R LM Immediate Recall, stories A and B; and BVMT-R Immediate Recall, trials 1-3); Delayed Recall (contributing tests: RAVLT Long-Delay Free Recall, WMS-R LM Delayed Recall, and BVMT-R Delayed Recall); and Executive Functioning (contributing tests: Trail Making Test Part B total time to completion, Stroop Neuropsychological Screening Test, color-word interference total items completed in 120 seconds, and WAIS-R Digit Symbol Coding total items completed in 90 seconds. The z-score for Trail Making Test Part B was multiplied by −1 before inclusion in the composite so that higher z-scores indicated better performance for all tests).

### Predictors of interest

At each study visit, participants completed a comprehensive neuropsychological testing battery, completed detailed medical history and lifestyle questionnaires, and provided blood samples, vitals, and other objective and physical assessments described elsewhere^9^. The three main lifestyle predictors—diet, physical activity, and cognitive activity—were based on self-report using the scales described below (see Table 1). We used data obtained from the first visit at which all three predictors were available (median = visit 5), and cognitive data from all visits from which the composites were available (range = visits 2-6).

**Table 1.** Description of self-reported health behaviors.

| Health Behavior | Self-Report Measure and Description |
| --- | --- |
| **Diet** | The Mediterranean-DASH Intervention for Neurodegenerative Delay Questionnaire (MIND; 15 items; Morris et al., 2015). The MIND diet score is based on consumption of 10 “brain healthy” food groups (e.g. leafy greens, nuts, berries, fish) and 5 unhealthy food groups (e.g., red meats, fried food, pastries/sweets). Range in WRAP: 3 – 14 (mean = 9.4, sd = 1.9) |
| **Physical Activity** | Participants completed the Women's Health Initiative physical activities questionnaire (McTieman et al., 2003). We calculated a total score of metabolic hours (MET) per week based on participants' responses to questions about the frequency and duration of walking outside the home for >10 minutes without stopping, and engagement in mild (e.g., golf), moderate (e.g., calisthenics), and vigorous (e.g., jogging) exercise<sup>18,41</sup>. For mild, moderate, and vigorous exercises, frequency ranged from none to $\geq 5$ days/week and duration ranged from <30 minutes to $\geq 1$ hour/session. Range in WRAP: 0 – 98 (mean = 18, sd = 15) |
| **Cognitive Activity** | The Florida Cognitive Activities Scale (FCAS) includes 25-items covering a range of cognitive activities (e.g., playing games, solving puzzles, cooking from new recipes) yielding a total cognitive activities score<sup>20</sup>. Range in WRAP sample: 14 – 70 (mean = 43.5, sd = 8.3) |

#### Physical activity

Participants answered questions about their physical activity based on a subset of items from the Women’s Health Initiative Observational study^16^. These questions were administered at visit 2 and each longitudinal visit thereafter. Details about this questionnaire are described elsewhere^17,18^; in brief, we constructed composite recreational physical activity variables based on several questions. Specifically, participants were asked about frequency, intensity, and duration of walking outside the home for more than 10 minutes; frequency and duration of strenuous exercise (examples provided included aerobics, jogging, swimming laps), moderate-intensity activities (e.g., biking, easy swimming, folk dancing), and low-intensity activities (slow-dancing, bowling, golf). We converted these responses to hours exercised per week, assigned metabolic equivalent (MET) values for each of the three intensity activities, and multiplied the MET level by hours exercised per week, as was done in previous work^17–19^, to obtain a variable “MET-hours per week.”

#### Cognitive activity

In order to quantify the type and frequency of engagement in cognitive activities, participants answered questions from the Florida Cognitive Activities Scale (FCAS)^20^. The FCAS is a 25-item scale that measures intensity, frequency and duration of a variety of activities differing in cognitive demand, including “playing chess, bridge or knowledge games,” “gardening,” and “preparing meals from new recipes.”^20^ Responses were weighted according to frequency and duration and summed to create the total “cognitive activities” variable.

#### Diet

To measure dietary behavior, participants completed a 15-item questionnaire capturing adherence to a hybrid of the Mediterranean-style diet and the Dietary Approaches to Stop Hypertension (DASH) diet termed Mediterranean-DASH Intervention for Neurodegenerative Delay (MIND)^4,6^. The MIND diet score is comprised of 10 brain healthy foods (e.g., natural, plant-based foods, berries and green leafy vegetables, whole grains, fish, olive oil) and 5 unhealthy food groups (red meats, butter/margarine, cheese, pastries/sweets, fried/fast food). To obtain a total MIND score, the frequency of consumption of each category was summed following guidelines published by Morris et al (2015)^6^; due to updates in the MIND questionnaire, coding varied slightly from the original publication (for details, see Table A in S1 Text for the revised MIND questionnaire and Table B in S1 Text for the coding decisions that differ from the original).

#### Other covariates

Covariates included age, sex, *APOE-ε4* risk score (a summative log-odds ratio of all *APOE* alleles^21^), and Wide Range Achievement Test – 3 (WRAT-3) Reading subtest standard score^22^. The WRAT-3 Reading subtest was used as a proxy for educational attainment, as previous literature suggests that reading level may be a more sensitive proxy for educational experience than self-reported years of education^23–25^.

### Data analysis

The FCAS and MIND questionnaires were added to the study after baseline; accordingly, health behavior measures were obtained from the first visit at which all health behavior data was available for a participant (median = visit 5). That is, the set of health behavior predictors are cross-sectional (time-invariant), while the cognitive outcome measures are longitudinal.

Given our goal of characterizing the combined influences of multiple health behavior predictors and related interactions, we opted to use an information theoretic (IT) modeling technique detailed by our group in previous work.^26^ In brief, the IT framework evaluates a set of plausible scientific hypotheses, and uses the relative strengths of information considered across all models to obtain model-averaged parameter estimates, instead of selecting a single model based on traditional model selection techniques. In so doing, after fitting all of the models of interest to each outcome (with differing combinations of health behavior predictors and interactions), results are combined across models in proportion to the relative strength of information each model contributes^26^. The IT framework also offers advantages over a traditional model selection approach (e.g., forward or backward selection), in that it allows comparison of fits across all models and reduces over-estimation of effect sizes compared to standard models.

#### The model set

Based on previous research indicating that physical activity, cognitive activity, and diet modified associations between time and cognition, we developed a set of 9 linear mixed-effects models to investigate in these analyses. Model 1 included the previously mentioned set of covariates; additional models built on Model 1 and incorporated various combinations of the 3 health behavior variables and interactions with age (the longitudinal covariate in our models). We used the age at each visit as the time variable in order to account for differences in baseline ages and time intervals between visits. We centered age at the average for ease of interpretation, with precision to two decimal places for improved temporal resolution.

For each outcome and each model in the set, we fit the model using a linear mixed effects structure (random effects were subject-specific intercepts). Health behavior predictors were converted to z-scores (mean=0, sd=1) for analyses. Further details about model fitting and diagnostics are detailed in our previous work^26^.

#### Model outputs

After fitting the models, model statistics were extracted, and the minimum Akaike’s Information Criterion-corrected (AICc) statistics were used to calculate the relative strength of information between each model and the best fitting model, which we then used to calculate model weights (smaller AICcs have larger weights). Regression coefficient results from each model were multiplied by their corresponding weights, and weighted results were summed together for the final model averaged result. Nonparametric bootstrapping of this process over 10,000 replicates was used to create 95% confidence intervals of the model averaged coefficients, using the quantile method; confidence intervals that contain zero are interpreted as non-significant. In addition to obtaining the model-averaged parameter estimates, we used the model log-likelihoods to conduct likelihood ratio tests of each of models 2-9 vs model 1 to test whether the addition of multiple behavior terms improved model fit over the covariates-only model. In exploratory/secondary results, we compare parameter estimates from the best fitting model with the model-averaged parameter estimates for an outcome only if the likelihood ratio tests for that model vs model 1 was significant at the α=0.05 level.

All analyses were performed in R version 3.4.0. Linear mixed effect regressions were fit using ‘lmer’ in the lme4 package; AICc-based model statistics were calculated using ‘aictab’ in the AICcmodavg package; bootstrapping was performed using HTCondor version 8.6.3.

## Results

### Sample characteristics

Sample characteristics at visit 2 (when all cognitive measures for the PACC were first available) are shown in Table 2. Mean(sd) age was 57.7(6.4) years, 67.6% were female, average years of education was 16.2(2.7) and 98.1% were non-Hispanic white.

**Table 2.** Sample demographic and health characteristics at Wave 2 data collection.

|  | n=828 |
| --- | --- |
| Age at first cognitive composite* (mean (sd)) | 57.7 (6.4) |
| Average years of Follow-up (mean (sd, median, range)) | 6.3 (sd=1.9, median=7, range=2-10) |
| Sex = Female (%) | 560 (67.6) |
| *APOE-ε4* Carrier (%) | 312 (37.7) |
| WRAT-3 Reading - Standard Score (mean (sd)) | 105.7 (9.3) |
| CES-D Depression Score (mean (sd)) | 6.6 (6.7) |
| Race (n, % non-Hispanic White) | 812 (98.1) |
| Years of Education (mean (sd)) | 16.2 (2.7) |
| Systolic Blood Pressure (mean (sd)) | 124.2 (15.9) |
| Diastolic Blood Pressure (mean (sd)) | 74.1 (9.6) |
| Total Cholesterol (mean (sd)) | 200.3 (37.7) |
| Body Mass Index (mean (sd)) | 28.7 (6.1) |
| Waist-Hip Ratio (mean (sd)) | 0.86 (0.10) |
| MMSE (mean (sd), median) | 29 (.9), 30 |
| AVLT Total (mean (sd)) | 52 (8) |
| Logical Memory Story Recall Delayed (mean (sd)) | 26 (6.9) |
| Digit Symbol Coding | 57 (10) |

Abbreviations: APOE Risk Score: a summative odds ratio of all APOE alleles (Darst et al., 2017); WRAT-3 – Wide Range Achievement Test (Wilkinson, 1984); CES-D – Center for Epidemiology Scale for Depression. MMSE: Mini-Mental State Examination (Folstein et al., 1975); AVLT: Rey-Auditory-Verbal Learning Test (Schmidt et al. 1996). \*The Wave 2 visit is “baseline” for cognition in these analyses as it was the first visit at which the analyzed composites could be calculated.

### IT framework results

Model fit statistics, differences between AICc and the best model’s AICc, model weights, log-likelihoods, and likelihood ratio tests are presented in Table 3 for each outcome. Model averaged coefficients and corresponding 95% confidence intervals for all non-intercept terms are depicted in Figures 1–4. Results are discussed by outcome below.

**Table 3.**
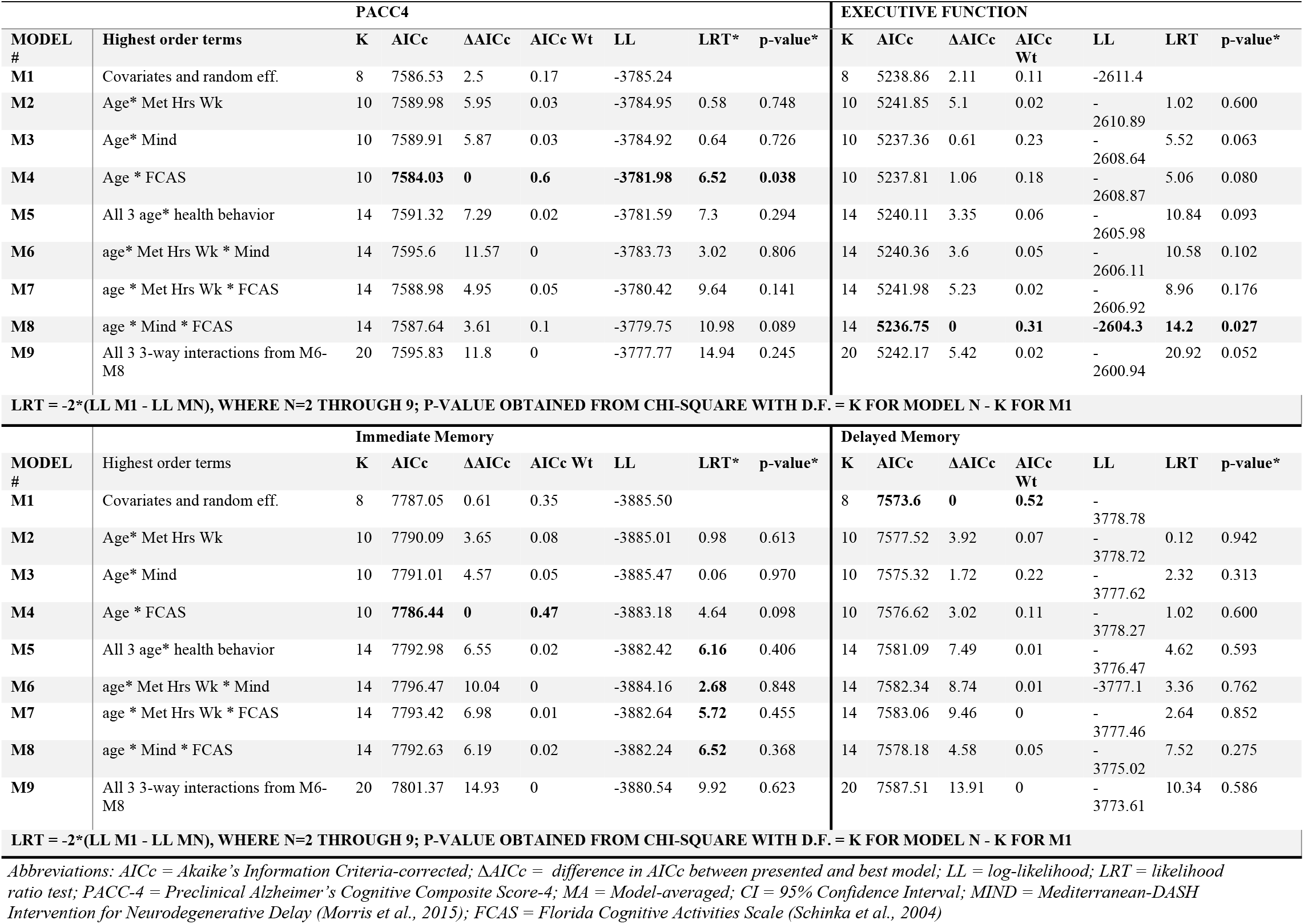
Model fit statistics, AICc, model weights, log-likelihood, and likelihood ratio tests

### Primary outcome: PACC4

The overall rate of change in PACC-4 z-scores for the sample, without adjusting for practice effects was .006(.25); the lowest quartile of change in PACC-4 was .30(.20) standardized score units per year.

The model-averaged estimates shown in Fig. 1 indicate that the main effects of age and WRAT-3 reading were significantly associated with the PACC4, such that participants with higher WRAT-3 reading and participants who were younger had better PACC4 scores. There were neither significant main effects nor significant interactions between age and any of the three modifiable behavior factors. The best fitting model of the set was model 4 (covariate model plus cognitive activity (FCAS) and cognitive activity(FCAS)*age interaction). The likelihood ratio test indicated that adding these terms resulted in a better fit than Model 1 (covariates-only model). Table 4 presents beta estimates and their confidence intervals for this model relative to the model-averaged parameter estimates. The FCAS*age model-averaged interaction beta and confidence interval were .006(−.002 – .02) compared with the model 4 beta and convidence interval of .007(.00003 − 0.01). Thus, although the model 4 beta and confidence interval did not contain zero, effects were weak, small, and similar to the IT model-averaged estimates.

**Fig 1.**
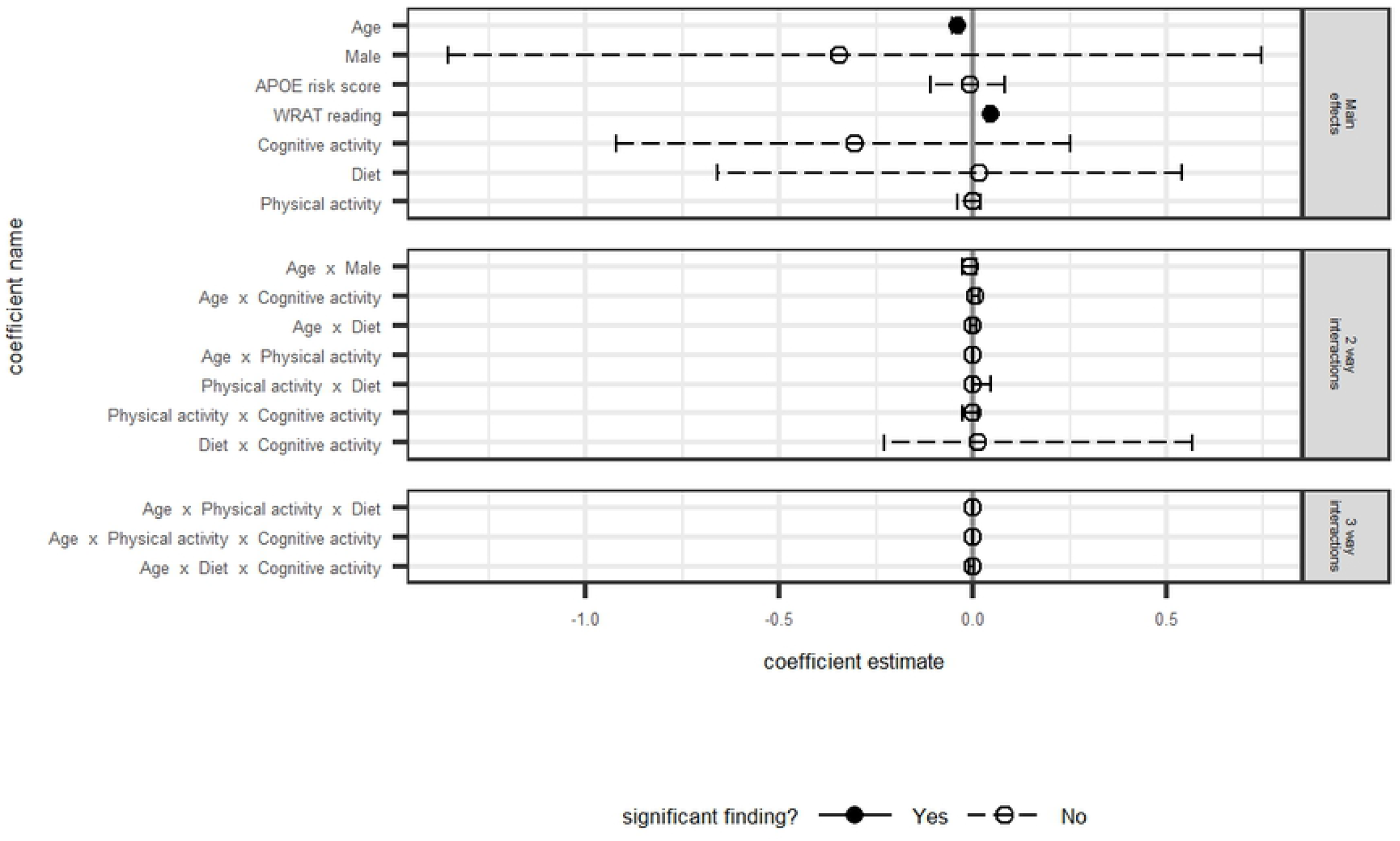
PACC: Model averaged regression coefficient estimates and 95% confidence intervals

**Table 4.**
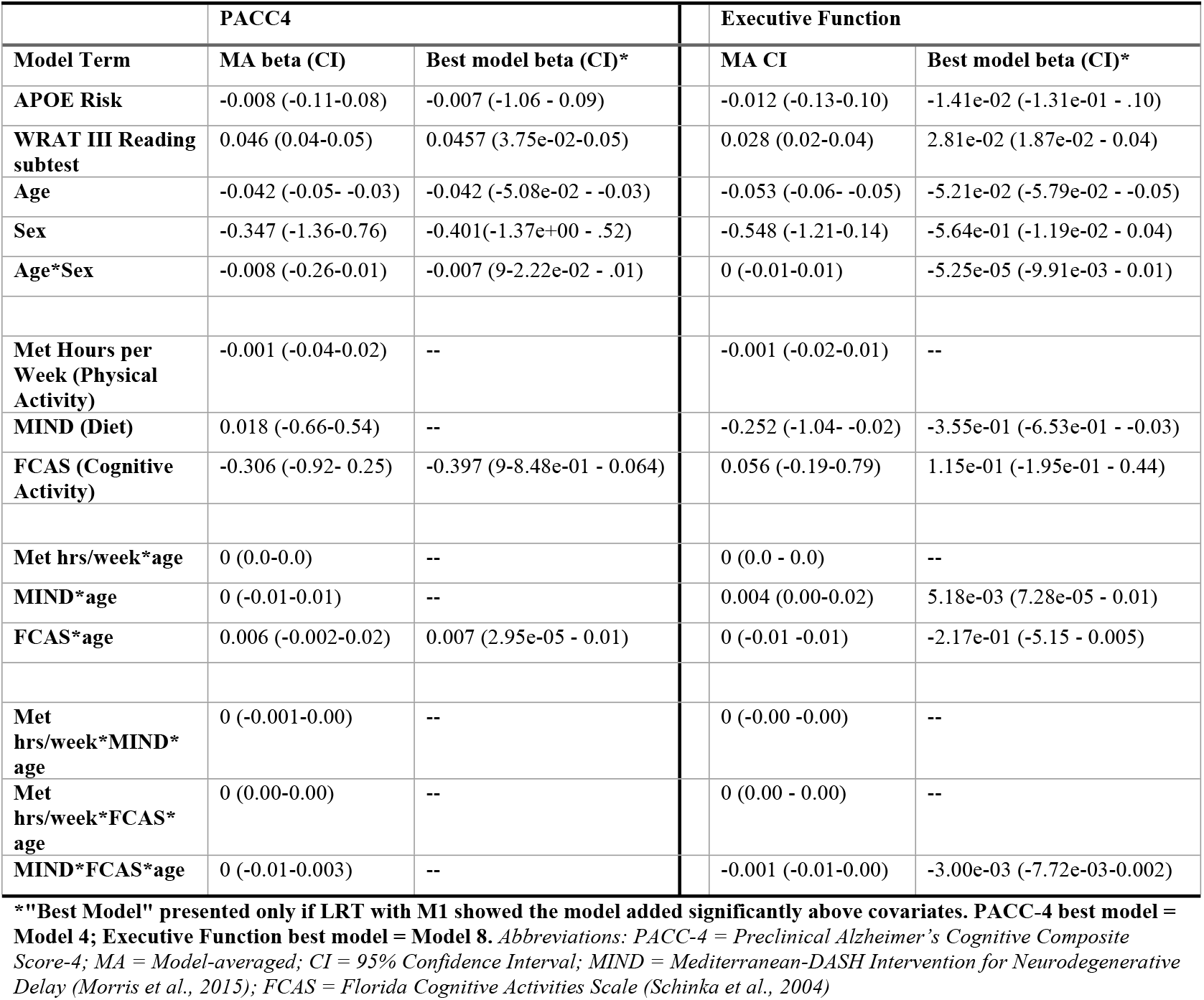
Beta estimates and 95% confidence intervals for best fitting models* relative to their model-averaged parameter estimates.

### Secondary outcomes

#### Immediate Memory composite

Main effects of age and WRAT-3 Reading were significantly associated with Immediate Memory, such that younger participants or those with higher WRAT-3 Reading scores performed better on Immediate Memory (Fig. 2). As with the PACC4, there were no significant main effects or interactions between the lifestyle behavior predictors of interest and the Immediate Memory outcome. The best fitting model of the set was model 4 (covariate model plus FCAS and FCAS*age interaction); the likelihood ratio test, however, was non-significant indicating that the addition of these terms did not fit the data better than the “non-health behaviors covariates only model” (i.e., Model 1).

**Fig 2.**
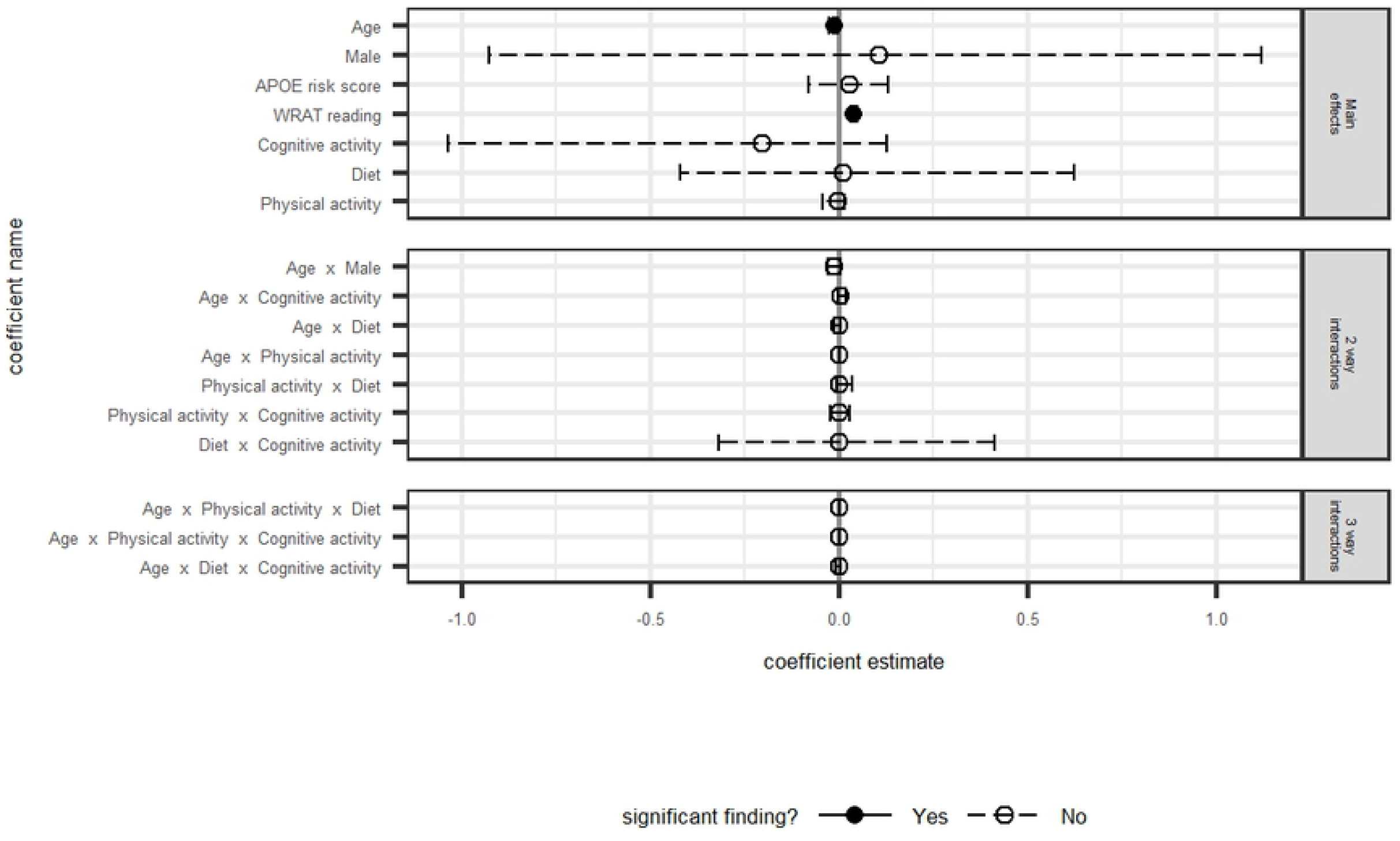
Immediate Memory: Model averaged regression coefficient estimates and 95% confidence intervals

#### Delayed Memory composite

None of the health behaviors were associated with delayed memory performance. The best fitting model of the set was model 1 (covariates only).

Higher WRAT-3 Reading scores were significantly associated with higher delayed memory scores. The significant interaction between age and sex indicated faster annual decline in men than in women (Fig. 3).

**Fig 3.**
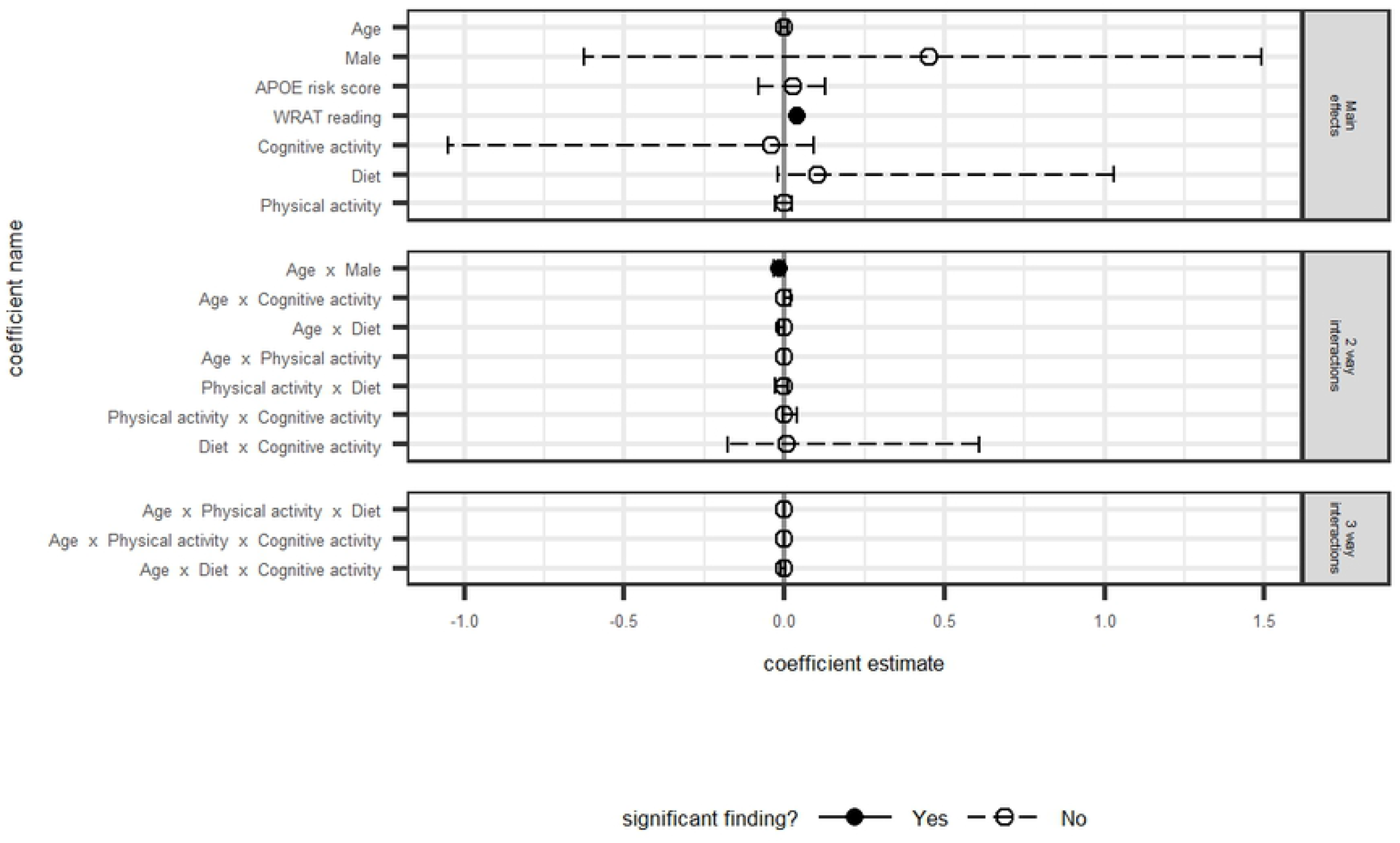
Delayed Memory: Model averaged regression coefficient estimates and 95% confidence intervals

#### Executive Function composite

The model averaged estimates shown in Fig. 4 indicate that main effects of age and WRAT-3 Reading were associated with executive functioning, such that younger participants and those with higher WRAT-3 scores performed better on the composite. A significant interaction for diet and age suggested that those participants who reported diets more congruent with the MIND diet showed slower decline over time on executive function (Fig. 5). No other predictors of interest were significantly associated with executive functioning. The best fitting model of the set was model 8 (covariate model plus MIND*FCAS*age interaction and corresponding lower-order terms). The likelihood ratio test indicated that this model fit the data better than Model 1. Results from this model were consistent with the model averaged estimates and indicated small effect sizes. Specifically, using model-averaged parameter estimatesand assuming FCAS is 0, those with MIND z-score = 1 had estimated annual decline of −.049 SDs per year vs −.057 SDs per year for those with MIND z-scores=−1. Using model 8 output, estimated annual change was −.047 vs − .057, respectively (for females with FCAS of 0). Table 4 presents beta estimates and their confidence intervals for this model relative to the model-averaged parameter estimates.

**Fig 4.**
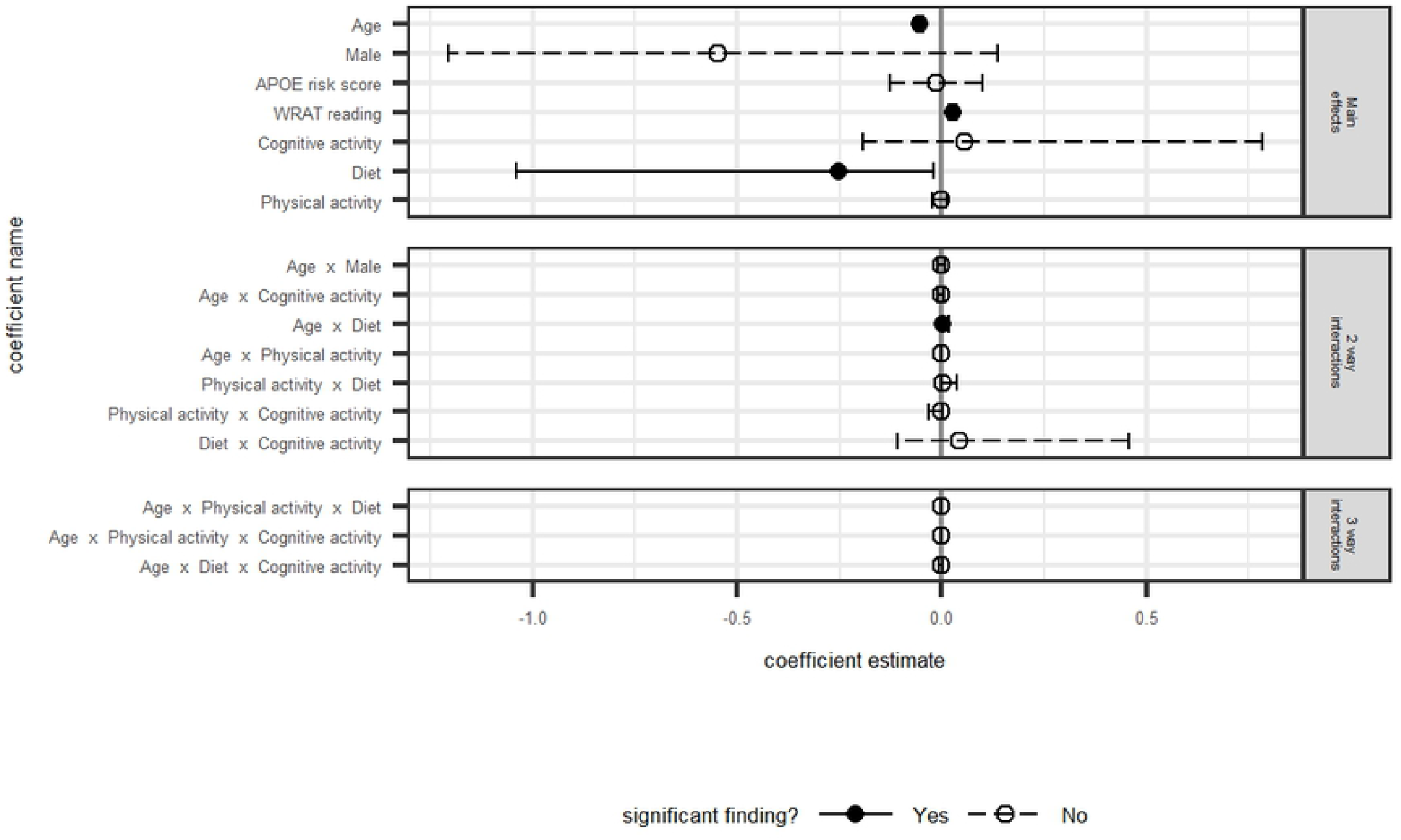
Executive Function: Model averaged regression coefficient estimates and 95% confidence intervals

**Fig 5.**
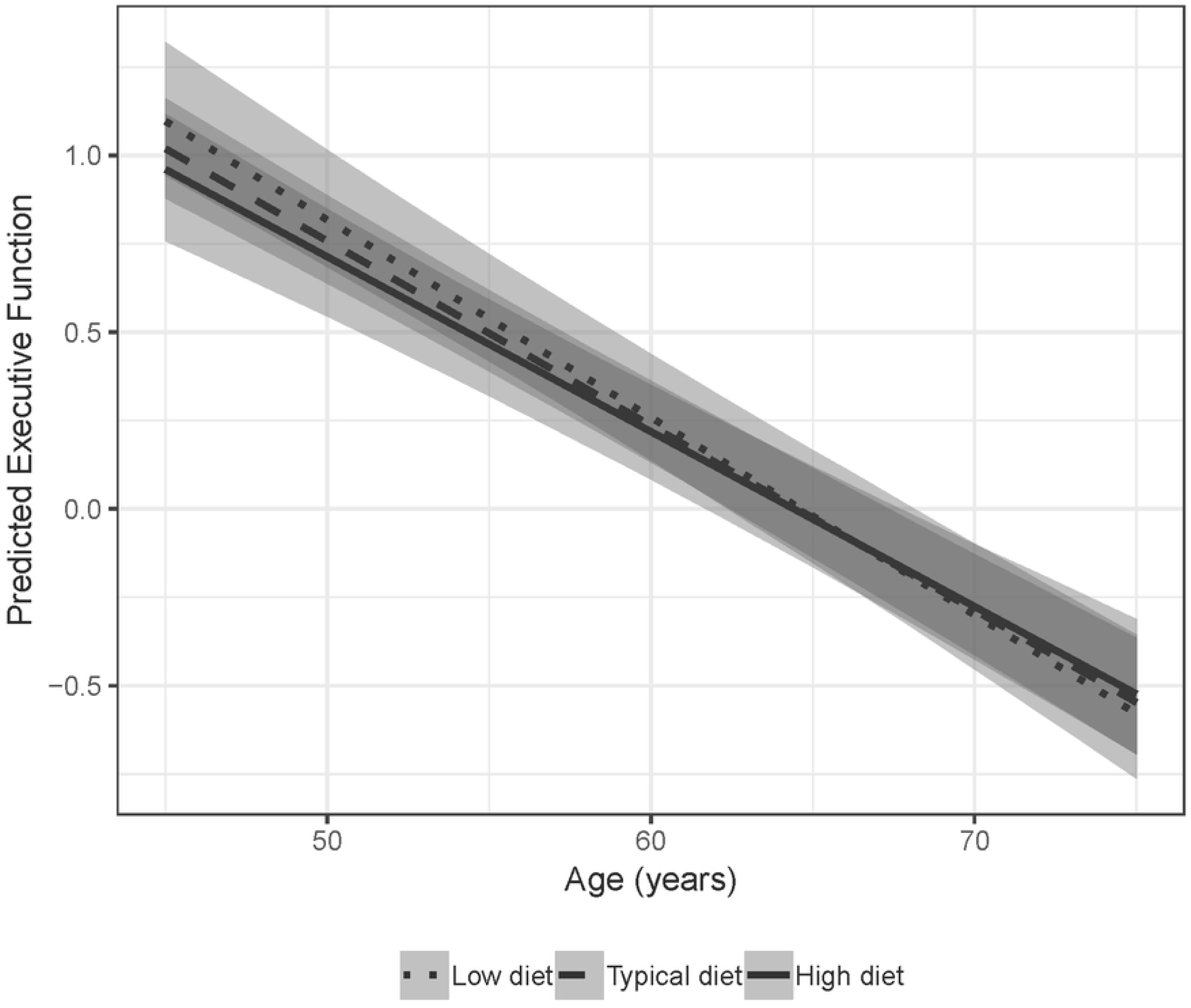
Model averaged predicted executive function performance by MIND diet with 95% confidence intervals

## Discussion

In this study, we examined whether multiple self-reported health behaviors – physical activity, cognitive activity, and diet – modified age-related cognitive trajectories in a group of cognitively unimpaired, late-middle-aged adults. General media narratives advise that such behaviors are protective, yet often the research behind these narratives has a singular focus (e.g., only diet or only physical activity). We used an information-theoretic model averaging approach so that we could include all three behaviors and multiple interactions at the same time, while reducing the risk of overfitting the data and at the same time effectively controlling for Type I error. We evaluated longitudinal outcomes in several cognitive domains that have been found to be sensitive to early cognitive decline, specifically; a global cognitive composite (Preclinical Alzheimer’s Cognitive Composite-4 [PACC4]^10^), immediate memory, delayed memory, and executive function). Our primary analysis failed to offer support that such behaviors influence cognitive decline in this cohort of late middle-aged people.

In this relatively large (n=828) and young sample (mean age at visit 5 follow-up was 65, standard deviation = 6.3) using a comprehensive approach, we found no associations between longitudinal cognition and self-reported diet or exercise. Analysis of secondary outcomes revealed a small effect regarding diet and executive function. Specifically, those participants who reported more adherence to a Mediterranean-DASH Intervention for Neurodegenerative Delay (MIND)^4^ diet showed slower decline on an executive function composite than those participants who reported less adherence to the MIND-style diet. The effect size for this interaction was quite small, despite the fact that several other studies have shown more robust associations between adherence to the MIND diet and reduced risk of cognitive decline and dementia^4–6^. One possible reason for this is the relatively young age of the cohort and corresponding reduced range of cognitive decline consistent with middle-agers; the average annual rate of change in cognitive composite z-scores was .005, while in the lowest quartile of change (i.e., a subset of participants potentially representing those with subclinical cognitive change) the mean change in PACC-4 was .30 (range = .13 to 1.40). Thus although the group as a whole is showing a small amount of change, there is a subgroup of individuals who are declining at a faster rate. Therefore it will be important to reexamine these associations as our sample ages and completes additional cognitive testing visits.

The lack of strong associations among health behaviors and longitudinal cognition in this late middle-aged cohort may also partially be explained by the fact that we did not consider objective measures of health (e.g., body mass index, blood pressure) or overall cardiovascular risk as covariates or mediating factors on cognition, as was done in other studies^27^. It is therefore possible that these health behaviors may be of little consequence in late middle age to young old age if chronic disease is not managed; future studies of individuals in late middle age that control for chronic disease and medical management of comorbidities, or examine cardiovascular risk as a mediating factor, will help provide a better understanding of these relationships. Second, we took a comprehensive approach to looking at health behaviors individually and when controlling for other health behaviors. Studies that have taken a similar approach have mixed results. For example, Sturman et al. (2005) examined whether participation in physical activity slowed cognitive decline after accounting for participation in cognitively stimulating activities; although physical activity was associated with a slower rate of cognitive decline, when adjusting for participation in cognitive activities, depression and vascular disease, the observed association of physical activity with to cognitive decline was no longer statistically significant^7^. Because individuals who engage in cognitive activity may be more likely to engage in physical activity and/or healthy diets, it is important to continue to attempt to elucidate the respective contributions that each behavior may have on cognition in late middle age and beyond^8^.

We used continuous measures of health behavior composite scores, similar to previous studies of these behaviors^28^. It is possible that categorical measures (e.g., tertiles indicating “high,” “medium”, and “low” engagement in behaviors) may provide added sensitivity, in that the discriminatory ability of these variable self-report measures to cognitive decline may lie in those who show the highest versus lowest engagement in behaviors^29^. Since it is likely that different health behaviors have an additive effect on health and possibly cognition, combining several health behaviors into an overall “health behavior index” may further elucidate these relationships. A recent large retrospective cohort study developed a weighted healthy lifestyle score, including no smoking, regular physical activity, healthy diet, and moderate alcohol consumption, which they categorized in to favoarable, intermediate and unfavorable lifestyles. Using this approach, the authors found that a favorable lifestyle was associated with lower dementia risk, including in those participants with higher genetic risk for dementia^30^.

It is important to consider the reliability of self-reported measures of lifestyle in a longitudinal cohort study such as WRAP. First, both the physical exercise composite and the MIND diet scores may either underestimate or overestimate actual engagement in these behaviors. The physical activity questions are broad (e.g., mild, moderate, strenuous) and participants may not have considered all possible sources of physical activity in their responses. Similarly, the 15-item revised MIND diet questionnaire had relatively few questions about food intake, including frequency of consumption of individual items, resulting in imprecise measurement of the food intake within each MIND diet component, therefore possibly underestimating the effect of diet on cognitive decline. Furthermore, some literature suggests that self-report may overestimate actual physical activity, possibly a result of effects such as social desirability bias^31^. The conflicting evidence about such measures suggests that self-reported health behaviors data should be interpreted with caution. Several large, multidomain intervention clinical trials are underway which will address the limitations of observational studies, including issues with self-report of health and behavior, but most of these studies are enrolling individuals who are 5-10 years older than the WRAP cohort, and still may not inform how lifestyle behaviors affect cognition in late middle-age. Examples of such trials include the Finnish Geriatric Intervention Study to Prevent Cognitive Impairment and Disability (FINGER) study, a multi-domain intervention study implementing dietary advice, a physical exercise program, cognitive training, and advice regarding management of metabolic and vascular risk factors^32^. One recent report showed that dietary improvement was associated with slower rate of decline in executive function (mean age of intervention and control groups = 69.5)^33^. Examples of other health behavior clinical trials aimed at reducing or preventing cognitive decline include Exercise in Adults with Mild Memory Problems (EXERT)^34^, Multidomain Alzheimer Preventive Trial (MAPT)^35^, and the U.S. Study to Protect Brain Health Through Lifestyle Intervention to Reduce Risk (POINTER)^36^.

Finally, it is possible that our sample is too young to see clinically meaningful associations between health behaviors and cognitive decline. For instance, while Morris et al. found that greater adherence to the MIND diet was associated with slower rates of decline in cognition^5^, this group was considerably older (mean age at follow-up = 81) than our WRAP sample (mean age at follow-up = 65). Similarly, some of the most compelling studies showing associations between health behaviors in midlife and reduced risk of cognitive decline either had dementia as the outcome^3–6^ or had older participant samples^3,28^. Furthermore, even though the cohort is enriched for *APOE-ε4* genotype (38% in this subset), our group has not found significant associations with the *APOE-ε4* allele and longitudinal cognition^9^, nor did we see any associations with *APOE-ε4* and cognition in the present study, despite the fact that APOE is a noted risk factor for cognitive decline^37^. This discrepancy may also be due to the younger age of the WRAP cohort and the reduced range of cognitive decline.

Our comprehensive information theoretic model averaging approach allowed us to test multiple predictors and multiple interactions, while avoiding issues common to model selection procedures (overfitting, anticonservative results, etc.). This methodological innovation lends credibility to our modest results. It should be noted that the WRAP sample is predominantly a self-selected family history cohort study, thereby enriched for AD-risk, and the majority of participants are highly educated, non-Hispanic white, and reside in the Upper Midwest, all of which may limit the generalizability of findings to the general population.

We did not examine whether or not these health behaviors had a protective effect on cognition in the context of disease (i.e., Alzheimer’s disease biomarkers including beta-amyloid plaques, phosphorylated tau, and neurodegeneration^38^), which will be an important area of future study in the WRAP cohort. In a 2015 study by Schultz and colleagues, higher cardiorespiratory fitness was found to modify the effects of amyloid burden on cognition, such that individuals with relatively high amyloid burden demonstrated better scores on cognitive testing if those individuals had high levels of cardiorespiratory fitness^39^. Additionally, Clark et al. (2019) found that hypertension and obesity moderated the relationship between cognitive decline and amyloid burden^40^. These results suggest that amyloid burden or other AD biomarkers may be an important variable to consider when assessing interactions between health behaviors and longitudinal cognitive trajectories in late middle-age.

Future directions for this work include: 1) examining multiple self-reported measures and the mediating effects of cardiovascular disease and other medical comorbitidies; 2) determining the effect of these behaviors on risk of Alzheimer’s disease by examining imaging and cerebrospinal fluid markers of beta-amyloid, phosphorylated tau, and neurodegeneration as outcomes; and 3) determining whether health behaviors moderate risk of cognitive decline or AD based on genetic markers of AD.

In conclusion, we found that self-reported physical activity, cognitive activity, and diet—when considered individually and together, but without inclusion of objective health measures—were not significantly associated with cognitive decline in a late middle-aged (spanning ages 45 to 79), cognitively unimpaired cohort at increased risk for AD. These findings may be important for public health messaging regarding health behaviors. It is possible that these behaviors alone may not be enough to reduce risk of cognitive decline in middle age, and that managing other modifiable risk factors such as hypertension or obesity, may be required for these behaviors to have an effect on cognition. Our future work including comorbidities, overall health behavior indices in the IT model averaging approach, and additional years of follow up, and associations with Alzheimer’s disease biomarkers, will help to address these questions further

## Acknowledgments

We gratefully acknowledge Allen Wenzel for creating the MIND Diet coding manual, the WRAP study team who have carefully acquired the longitudinal data, and the WRAP participants who make this research possible. The authors of this manuscript have no conflicts of interest to report.

## Supporting information

**S1 Table. Adapted MIND diet questionnaire* used in the WRAP study**. *Adapted from Morris, M. C., Tangney, C. C., Wang, Y., Sacks, F. M., Bennett, D. A., & Aggarwal, N. T. (2015). MIND diet associated with reduced incidence of Alzheimer’s disease. *Alzheimer’s & Dementia*, 11(9), 1007-1014.

**S2 Table. Comparison of coding schemes between Morris et al. (2015) and that used in WRAP**. * Morris, M. C., Tangney, C. C., Wang, Y., Sacks, F. M., Barnes, L. L., Bennett, D. A., & Aggarwal, N. T. (2015). MIND diet slows cognitive decline with aging. *Alzheimer’s & dementia*, 11(9), 1015-1022. †Responses were converted to real numbers where possible. If the conversion didn’t translate to a real number, or if the resulting number was less than zero, the response was treated as a missing value. It should also be noted that if the participant responded with “>” a number, a value of “.2” was added to the converted value, i.e. a field value of “> 1” was assigned a value of “1.2”. Likewise, if a “<” was found at the beginning of the field, a value of “.2” was subtracted from the converted value, i.e. a field value of “< 1” was assigned a value of “.8”. Any missing field values were not included in the “MIND Diet Score” calculation. For those participants with missing values, a linear “extrapolated sum” was computed based on the number of existing scores and total number of items. For example, if there were three responses missing, and the sum of all scores present was 6.5, then the extrapolated sum would be calculated as follows: (6.5 / 12) * 15 = 8.125.

**This coding scheme was implemented due to participant reports that did not fall within any of the three specified ranges of the published criteria (Morris et al., 2015).

## References

1. 2018 Alzheimer’s disease facts and figures. Alzheimer’s & Dementia. 2018;14(3):367–429. doi:10.1016/j.jalz.2018.02.001

2. Fratiglioni L, Paillard-Borg S, Winblad B. An active and socially integrated lifestyle in late life might protect against dementia. The Lancet Neurology. 2004;3(6):343–353.

3. Paillard-Borg S, Fratiglioni L, Xu W, Winblad B, Wang H-X. An active lifestyle postpones dementia onset by more than one year in very old adults. Journal of Alzheimer’s Disease. 2012;31(4):835–842.

4. Morris MC, Tangney CC, Wang Y, Barnes LL, Bennett D, Aggarwal N. MIND diet score more predictive than DASH or Mediterranean diet scores. Alzheimer’s & Dementia: The Journal of the Alzheimer’s Association. 2014;10(4):P166.

5. Solfrizzi V, Custodero C, Lozupone M, et al. Relationships of dietary patterns, foods, and micro- and macronutrients with Alzheimer’s disease and late-life cognitive disorders: A systematic review. Journal of Alzheimer’s Disease. 2017;59(3):815–849.

6. Morris MC, Tangney CC, Wang Y, Sacks FM, Bennett DA, Aggarwal NT. MIND diet associated with reduced incidence of Alzheimer’s disease. Alzheimer’s & Dementia. 2015;11(9):1007–1014.

7. Sturman MT, Morris MC, Leon CFM de, Bienias JL, Wilson RS, Evans DA. Physical Activity, Cognitive Activity, and Cognitive Decline in a Biracial Community Population. Arch Neurol. 2005;62(11):1750–1754. doi:10.1001/archneur.62.11.1750

8. Spring B, Moller AC, Coons MJ. Multiple health behaviours: overview and implications. J Public Health (Oxf). 2012;34(Suppl 1):i3–i10. doi:10.1093/pubmed/fdr111

9. Johnson SC, Koscik RL, Jonaitis EM, et al. The Wisconsin Registry for Alzheimer’s Prevention: A review of findings and current directions. Alzheimer’s & Dementia: Diagnosis, Assessment & Disease Monitoring. 2018;10:130–142. doi:10.1016/j.dadm.2017.11.007

10. Donohue MC, Sperling RA, Salmon DP, et al. The preclinical Alzheimer cognitive composite: measuring amyloid-related decline. JAMA neurology. 2014;71(8):961–970.

11. Schmidt M. Rey Auditory Verbal Learning Test: RAVLT: A Handbook. Los Angeles, CA: Western Psychological Services; 1996.

12. Wechsler D. WMS-R: Wechsler Memory Scale--Revised: Manual. San Antonio: Psychological Corp.: Harcourt Brace Jovanovich; 1987.

13. Wechsler D. WAIS-R, Wechsler Adult Intelligence Scale-Revised, Manual. San Antonio, TX: The Psychological Corporation; 1981.

14. Folstein MF, Folstein SE, McHugh PR: Mini-Mental State: A practical method for grading the cognitive state of patients for the clinician. J Psychiatr Res 1975;12:189–198.

15. Clark LR, Racine AM, Koscik RL, et al. Beta-amyloid and cognitive decline in late middle age: Findings from the WRAP study. Alzheimers Dement. 2016;12(7):805–814. doi:10.1016/j.jalz.2015.12.009

16. Design of the Women’s Health Initiative Clinical Trial and Observational Study. Controlled Clinical Trials. 1998;19(1):61–109. doi:10.1016/S0197-2456(97)00078-0

17. Head D, Bugg JM, Goate AM, et al. Exercise Engagement as a Moderator of the Effects of APOE Genotype on Amyloid Deposition. Arch Neurol. 2012;69(5):636–643. doi:10.1001/archneurol.2011.845

18. Schultz SA, Boots EA, Almeida RP, et al. Cardiorespiratory fitness attenuates the influence of amyloid on cognition. J Int Neuropsychol Soc. 2015;21(10):841–850. doi:10.1017/S1355617715000843

19. Boots EA, Schultz SA, Oh JM, et al. Cardiorespiratory fitness is associated with brain structure, cognition, and mood in a middle-aged cohort at risk for Alzheimer’s disease. Brain Imaging and Behavior. 2015;9(3):639–649. doi:10.1007/s11682-014-9325-9

20. Schinka JA, Mcbride A, Vanderploeg RD, Tennyson K, Borenstein AR, Mortimer JA. Florida Cognitive Activities Scale: initial development and validation. Journal of the International Neuropsychological Society. 2005;11(1):108–116.

21. Darst BF, Koscik RL, Racine AM, et al. Pathway-specific polygenic risk scores as predictors of ß-amyloid deposition and cognitive function in a sample at increased risk for Alzheimer’s disease. J Alzheimers Dis. 2017;55(2):473–484. doi:10.3233/JAD-160195

22. Wilkinson GS. WRAT-3: Wide Range Achievement Test. Wide Range; 1993.

23. Manly JJ. Deconstructing race and ethnicity: implications for measurement of health outcomes. Medical care. 2006;44(11):S10–S16.

24. Sisco S, Gross AL, Shih RA, et al. The Role of Early-Life Educational Quality and Literacy in Explaining Racial Disparities in Cognition in Late Life. J Gerontol B Psychol Sci Soc Sci. 2015;70(4):557–567. doi:10.1093/geronb/gbt133

25. Manly JJ, Byrd DA, Touradji P, Stern Y. Acculturation, Reading Level, and Neuropsychological Test Performance Among African American Elders. Applied Neuropsychology. 2004;11(1):37–46. doi:10.1207/s15324826an1101_5

26. Koscik RL, Norton DL, Allison SL, et al. Characterizing the Effects of Sex, APOE ε4, and Literacy on Mid-life Cognitive Trajectories: Application of Information-Theoretic Model Averaging and Multi-model Inference Techniques to the Wisconsin Registry for Alzheimer’s Prevention Study. J Int Neuropsychol Soc. December 2018:1–15. doi:10.1017/S1355617718000954

27. Morris MC, Tangney CC, Wang Y, et al. MIND diet slows cognitive decline with aging. Alzheimer’s & dementia. 2015;11(9): 1015–1022.

28. Wilson RS, Leon CFM de, Barnes LL, et al. Participation in Cognitively Stimulating Activities and Risk of Incident Alzheimer Disease. JAMA. 2002;287(6):742–748. doi:10.1001/jama.287.6.742

29. Scarmeas N, Stern Y. Cognitive reserve and lifestyle. Journal of clinical and experimental neuropsychology. 2003;25(5):625–633.

30. Lourida I, Hannon E, Littlejohns TJ, et al. Association of Lifestyle and Genetic Risk With Incidence of Dementia. JAMA. July 2019. doi:10.1001/jama.2019.9879

31. Brenner PS, DeLamater JD. Social Desirability Bias in Self-reports of Physical Activity: Is an Exercise Identity the Culprit? Soc Indic Res. 2014;117(2):489–504. doi:10.1007/s11205-013-0359-y

32. Ngandu T, Lehtisalo J, Solomon A, et al. A 2 year multidomain intervention of diet, exercise, cognitive training, and vascular risk monitoring versus control to prevent cognitive decline in at-risk elderly people (FINGER): a randomised controlled trial. Lancet. 2015;385(9984):2255–2263. doi:10.1016/S0140-6736(15)60461-5

33. Lehtisalo J, Levälahti E, Lindström J, et al. Dietary changes and cognition over 2 years within a multidomain intervention trial—The Finnish Geriatric Intervention Study to Prevent Cognitive Impairment and Disability (FINGER). Alzheimer’s & Dementia. 2019;15(3):410–417. doi:10.1016/j.jalz.2018.10.001

34. Baker LD, Cotman C, Morrison RH, et al. EXERT: A PHASE 3 MULTI-SITE RANDOMIZED CONTROLLED TRIAL OF AEROBIC EXERCISE IN MCI — STUDY DESIGN AND METHODS. Alzheimer’s & Dementia: The Journal of the Alzheimer’s Association. 2017;13(7):P613. doi:10.1016/j.jalz.2017.06.672

35. Vellas B, Carrie I, Gillette-Guyonnet S, et al. MAPT STUDY: A MULTIDOMAIN APPROACH FOR PREVENTING ALZHEIMER’S DISEASE: DESIGN AND BASELINE DATA. J Prev Alzheimers Dis. 2014;1(1):13–22.

36. U.S. Study to Protect Brain Health Through Lifestyle Intervention to Reduce Risk – Full Text View – ClinicalTrials.gov. https://clinicaltrials.gov/ct2/show/NCT03688126. Accessed July 29, 2019.

37. Howieson DB, Camicioli R, Quinn J, et al. Natural history of cognitive decline in the old old. Neurology. 2003;60(9):1489–1494. doi:10.1212/01.wnl.0000063317.44167.5c

38. Jack CRJ, Bennett DA, Blennow K, Carrillo MC, Dunn B, Elliott C. NIA-AA Research Framework: Towards a Biological Definition of Alzheimer’s Disease.; 2017.

39. Schultz SA, Boots EA, Almeida RP, et al. Cardiorespiratory fitness attenuates the influence of amyloid on cognition. J Int Neuropsychol Soc. 2015;21(10):841–850. doi:10.1017/S1355617715000843

40. Clark LR, Koscik RL, Allison SL, et al. Hypertension and obesity moderate the relationship between ß-amyloid and cognitive decline in midlife. Alzheimers Dement. 2019;15(3):418–428. doi:10.1016/j.jalz.2018.09.008

